# Human neuroblastoma SH-SY5Y cells treated with okadaic acid express phosphorylated high molecular weight tau-immunoreactive protein species

**DOI:** 10.1101/284265

**Authors:** Mirta Boban, Terezija Miškić, Mirjana Babić Leko, Patrick R. Hof, Goran Šimić

## Abstract

Recent data suggest that early stages of Alzheimer’s disease (AD) are characterized by an abnormally high phosphorylation of microtubule-associated protein tau and truncation of its C-terminus. Tau hyperphosphorylation may result from the downregulation of phosphatases, especially protein phosphatase 2A. In an attempt to model and analyze these molecular events we treated SH-SY5Y cells with okadaic acid (OA), an inhibitor of protein phosphatases. In addition to the low molecular weight tau, such treatment lead to the appearance of heat-stable protein species with apparent high molecular weight around 100 kDa, which were immunoreactive to tau antibodies against phosphorylated Ser202 and phosphorylated Ser396. Based on the observation that high molecular weight tau-immunoreactive proteins (HMW-TIP) correspond to the predicted size of two monomers of tau, one possibility is that HMW-TIP represent tau oligomers. The absence of HMW-TIP detection by anti-total tau antibodies used may be a consequence of epitope masking, or a combination of epitope masking and protein truncation. We noted the stability of HMW-TIP in the presence of strong denaturing agents, such as urea and guanidine, as well as upon partial dephosphorylation by the alkaline phosphatase. Moreover, as HMW-TIP did not dissociate the presence of β-mercaptoethanol, it was also independent from disulfide bonds. Taken together, these data show that OA treatment of SH-SY5Y cells induces the appearance of HMW-TIP, which may represent tau oligomer or tau-crossreactive phospho-proteins. Our findings have implications for further studies of tau under the conditions of protein phosphatase downregulation.

## 1. Introduction

One of the neuropathological hallmarks of Alzheimer’s disease (AD) is the aggregation of tau protein in neurofibrillary tangles (NFT) in the neurons and glia cells in the brain. Tau (tubulin-associated unit) is a microtubule-associated protein, expressed mainly in neurons of the central nervous system [1], where its primary function is to modulate microtubule dynamics for maintaining axonal cytoskeleton. In adult human brain, there are six tau splicing isoforms ranging from 352 to 441 amino acids (for review, see [2]). Isoforms differ by the absence or presence of one or two 29 amino acid inserts in the N-terminal part in combination with either three or four microtubule-binding (MTB) repeat regions in the C-terminal part (for review, see [3]).

Tau is normally a natively unfolded protein [4] with little tendency for aggregation, therefore an intriguing but largely unresolved question is to understand what causes wild-type tau molecules to oligomerize and subsequently form larger aggregates that form NFT. Pathways leading to tau aggregation are largely unclear. However, tau propensity to aggregate may be driven by posttranslational modifications, mainly phosphorylation and C-terminal truncation [5]. Despite a paucity of stable secondary structure, tau in solution seems to form a transient global fold in which N- and C-terminal arms fold over the central MTB repeat domain that contains the sequences with a tendency to form β-sheets and aggregate (known as the “paper clip” model [6]).

Phosphorylation decreases affinity of tau for microtubules and makes it more prone to aggregation (for review, see [7]). In AD brains more than thirty tau amino acid residues are specifically phosphorylated, including Ser202 and Ser396 [8]. Hyperphosphorylation of tau in AD seems to be a consequence of a disrupted balance between kinase and phosphatase activity. Several protein phosphatases were found dysregulated in AD brain, including PP2A, PP1 and PP5, which are all capable of dephosphorylating tau *in vitro* (for review, see [9]. The main regulator of tau dephosphorylation in healthy human brain appears to be PP2A, since its activity accounted for about 70% of the tau dephosphorylation in the assay using brain extracts [10]. PP2A has been shown to dephosphorylate tau *in vitro* at multiple sites, including Ser202 and Ser396 [11]. PP2A affects tau phosphorylation levels not only directly, but also indirectly by regulating the activities of several tau kinases, most notably glycogen synthase kinase 3β (GSK-3β) and Ca^2+^/calmodulin-dependent protein kinase II (CaMKII) [12].

To investigate the effects of protein phosphatase downregulation on the process of tau phosphorylation and aggregation, we took an approach of protein phosphatase inhibition based on okadaic acid (OA) application to cultured cells. OA is a cell-permeable potent inhibitor of PP2A and PP4, and of PP5 and PP1 [13]. As a cell culture model we chose human neuroblastoma cell line SH-SY5Y. These cells differentiate into neuron-like cells that have the endogenous expression of tau [14-16]. We observed that the incubation of SH-SY5Y cells with OA leads to the appearance of high molecular weight protein species immunoreactive to tau antibodies against phosphorylated Ser202 and phosphorylated Ser396.

## 4. Materials and Methods

### 4.1. Cell culture

Cells SH-SY5Y (ECACC, 94030304) were grown in Dulbecco’s modified Eagle medium (DMEM, Gibco, Gaithersburg, MD, USA, cat. no. 31885-049) supplemented with 10% fetal bovine serum (FBS, Gibco, cat. no. 10270106), 1% L-glutamine (Gibco, cat. no. 25030024), 1% non-essential amino acids (Sigma-Aldrich, Darmstadt, Germany, cat. no. M7145) and 1% penicillin-streptomycin (Gibco, cat. no. 15140-122) at 37°C in humidified atmosphere with 5% CO_2_. Unless indicated otherwise, undifferentiated SH-SY5Y cells grown to 60-90% confluency were used. For differentiation into neuron-like type, we followed the protocol described by Encinas and colleagues [17] with minor modifications. Briefly, cells were seeded at density of 10,000 cells/cm^2^ in the medium described above. The following day a medium containing 10 μM all-trans retinoic acid was added to the cells and incubated for five days, the medium being replaced every other day. Cells were washed with serum-free medium and incubated in medium containing 1% FBS and 50 ng/ml brain-derived neurotrophic factor (BDNF, Sigma-Aldrich, cat. no. SRP3014) for two days. Cells were photographed using phase contrast microscopy (Zeiss, Oberkochen, Germany). Where indicated, cells were treated with 100 or 150 nM OA (Abcam, Cambridge, UK, cat. no. ab120375) added from a concentrated solution in dimethyl sulfoxide (DMSO). DMSO was used as a negative control.

### 4.2. Preparation of cell lysates and protein extraction

Upon harvesting, cells were washed in ice-cold Tris-buffered saline (TBS) buffer (20 mM Tris-HCl pH 7.4, 150 mM NaCl) and resuspended in Laemmli buffer (50 mM Tris-HCl pH 6.8, 2% sodium dodecyl sulfate [SDS], 2 mM ethylenediaminetetraacetic acid [EDTA], 10% glycerol, 1 “Complete Mini” protease inhibitor cocktail tablet per 10 ml [Roche Diagnostics, Basel, Switzerland cat. no. 11836170001, Roche Diagnostics, Basel, Switzerland], 1 mM phenylmethylsulfonyl fluoride, 10 mM sodium fluoride, 20 mM β-glycerophosphate, and 2 mM sodium orthovanadate). For experiments shown in Figures 1C, 2B and 3A cells were resuspended in 2x Laemmli buffer without prior washing step. Unless otherwise indicated, 2-5% β-mercaptoethanol was present in the lysis buffer and samples were denatured at 90°C or 95°C for 5-15 min. For the experiment shown in Figures 1A and 1B, cells were resuspended in ice-cold radioimmunoprecipitation assay (RIPA) buffer (20 mM Tris-HCl pH 7.4, 1% Triton X-100, 0.1 % sodium dodecyl sulfate, 0.5 % sodium-deoxycholate, 150 mM NaCl, 1 mM EDTA, 10 % glycerol, protease and phosphatase inhibitors as described above), drawn through 23G needle 8 times and incubated on ice for 30 min. Samples were centrifuged at 14,000 g for 10 min at 4°C, Laemmli buffer containing 5% β-mercaptoethanol was added to the supernatant and incubated at 95°C for 5 min. For urea treatment, cells were resuspended in 2x Laemmli buffer containing 8 M urea, samples were incubated at 65 C° for 10 min, centrifuged at 20,000 g for 20 min at room temperature (RT) and the supernatant was collected.

**Figure 1.**
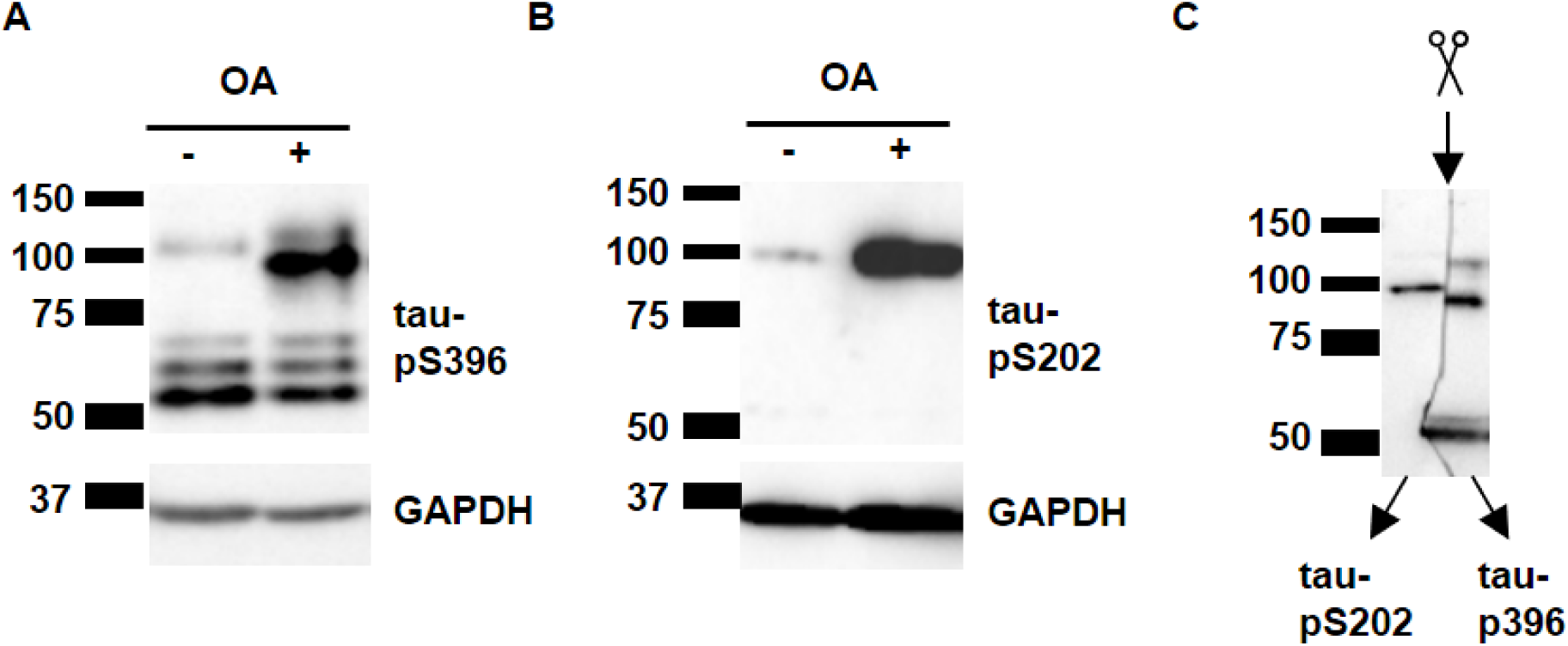
OA treatment of SH-SY5Y cells induces expression of a high molecular weight protein species immunoreactive to phospho-tau. Cells were treated with 150 nM (A, B) or 100 nM (C) OA for two hours. Total cells lysates were analyzed by immunoblotting using antibodies that detect tau-pS396 and tau-pS202 (CP13). GAPDH served as a loading control (A, B). Note a strong band migrating around 100-kDa in lysates of OA-treated cells. (C) Nitrocellulose membrane with transferred proteins was cut vertically in the middle of the lane (scissors) and the halves were incubated with CP13 or anti-tau-pS396 antibodies. Membranes were placed side-by-side and imaged simultaneously.

### 4.3. Cell extraction by guanidine-hydrochloride

Cells harvested by trypsinization were washed in ice-cold phosphate-buffered saline (PBS) and resuspended in a solution containing 6 M guanidine-hydrochloride, 20 mM Tris-HCl pH 7.4 and 100 mM NaCl, incubated at RT for 10 min. Suspension was drawn through 23 G needle and incubated at RT for 20 min, followed by centrifugation at 20,000 g for 20 min at RT. The supernatant was collected and the centrifugation step repeated. Guanidine hydrochloride forms a precipitate with SDS and is thus incompatible with SDS-PAGE [18]. To remove guanidine hydrochloride, proteins from the supernatant were precipitated by incubation in 90% ethanol at −20° C for 1 h. The samples were centrifuged at 20,000 g for 20 min at 4°C, the pellet was washed in absolute ethanol and air-dried. Precipitated proteins were solubilized in 2x Laemmli buffer (100 mM Tris-HCl pH 6.8, 4% SDS, 4 mM EDTA, 10% glycerol, 1 Roche “Complete Mini” protease inhibitor cocktail tablet per 10 ml, 1 mM phenylmethylsulfonyl fluoride, 10 mM sodium fluoride, 20 mM β-glycerophosphate and 2 mM sodium orthovanadate) containing 8 M urea, incubated at 37°C for 20 min, and centrifuged at 20,000 g for 20 min at RT. β-mercaptoethanol was added to supernatant to a final concentration of 5% and the centrifugation step repeated.

### 4.4. Heat-stable protein fraction

To prepare heat stable fraction, we followed the protocol described by Petry *et al.* with some modifications [19]. Briefly, the cells harvested by Versene (Gibco, cat. no. 15040-066) were washed in PBS buffer, resuspended in 250 μl lysis buffer (50 mM Tris-HCl pH 7.4, 1% Triton X-100, 0.25 % sodium-deoxycholate, 150 mM NaCl, 1 mM EDTA, 10% glycerol, 1 Roche “Complete Mini” protease inhibitor cocktail tablet per 10 ml, 1 mM phenylmethylsulfonyl fluoride, 10 mM sodium fluoride, 20 mM β-glycerophosphate, and 2 mM sodium orthovanadate), incubated on ice for 10 min, drawn through 23 G needle eight times, incubated on ice for 20 min and centrifuged at 20,000 g for 10 min at 4°C. The supernatant was collected and incubated at 95°C for 10 min, followed by centrifugation at 20,000 g for 20 min at 4°C. Supernatant containing the heat-stable fraction was separated from the pellet. Approximately 160 μl of supernatant was incubated with one third of the volume of concentrated Laemmli buffer (8% SDS, 200 mM Tris-HCl pH 6.8, 8 mM EDTA, 20% glycerol). The pellet was resuspended in a similar volume of radioimmunoprecipitation assay (RIPA) buffer and concentrated Laemmli buffer.

### 4.5. Alkaline phosphatase treatment of protein lysates

The cells were treated with 100 nM OA for 2 h, trypsinized, washed in PBS buffer and resuspended in lysis buffer (50 mM Tris-HCl pH 7.9, 1% Triton X-100, 0.25% sodium-deoxycholate, 150 mM NaCl, 1 mM MgCl^2^, 1 Roche “Complete Mini” protease inhibitor cocktail tablet per 10 ml, 1 mM phenylmethylsulfonyl fluoride, 10 mM sodium fluoride, 20 mM β-glycerophosphate), incubated on ice for 10 min, drawn through 23 G needle eight times, incubated on ice for 20 min followed by 5 min centrifugation at 20,000 g at 4°C. The supernatant was incubated at 95°C for 10 min, followed by centrifugation at 20,000 g for 20 min at 4°C. The lysate (75 μl) was incubated with 25 U of alkaline phosphatase (Promega, Fitchburg, WI, USA, cat. no. M2825) at 37 C° for 30 min, followed by addition of Laemmli buffer and 5 min incubation at 95°C.

### 4.6. Immunoblotting

Proteins in cell extracts were separated on 7.5 % or 10% SDS-PAGE gels and transferred onto a nitrocellulose membrane. The membrane was blocked in 5% milk diluted in TBS or phosphate-buffered saline (PBS; 137 mM NaCl, 2.7 mM KCl, 10 mM Na_2_HPO_4_, 1.8 mM KH_2_PO_4_) and incubated with primary antibody diluted in TBS or PBS containing 5% milk or 3% bovine serum albumin, overnight at 4°C. The primary antibodies and their working dilutions used were: anti-tau-pS396 (Abcam, cat. no. ab109390, 1:10,000), Tau46 (Sigma-Aldrich, T9450, 1:1,000), anti-tau-pS202 (CP13, mouse monoclonal, gift of Prof. Peter Davies, 1:500), anti-total tau mouse monoclonal antibody CP27 (gift of Prof. Peter Davies, 1:250), Tau13 (Santa Cruz Biotechnology, Santa Cruz, CA, USA, cat. no. sc-21796, 1:250) anti-MAP2 (clone HM-2, Sigma-Aldrich, cat. no. M4403, 1:1,000) and anti-GAPDH (glyceraldehyde 3-phosphate dehydrogenase, Cell Signaling, Leiden, Netherlands, cat. no. 2118, 1:1000 to 1:2,000). The secondary antibodies were horse radish peroxidase-linked IgG (Cell Signaling, cat. no. 7074 and 7076). Proteins were visualized using Super Signal West substrate (Thermo Scientific, Waltham, MA, USA, Pico cat. no. 34077 or Femto cat. no. 34095) and imager ChemiDoc XRS+ with ImageLab software (BioRad Laboratories, Hercules, CA, USA).

## 2. Results

### 2.1. OA treatment of SH-SY5Y cells induced expression of 100 kDa proteins immunoreactive with anti-tau-pS202 and anti-tau-pS396 antibodies

To inhibit the activity of protein phosphatases, we incubated undifferentiated neuroblastoma SH-SY5Y cells that endogenously express tau with 100 nM OA for two hours. Proteins from total cell protein lysates were separated by denaturing SDS polyacrylamide gel electrophoresis (SDS-PAGE). Tau phosphorylation at specific epitopes was assessed by immunoblot using phospho-specific tau antibodies. Using the antibody that recognizes tau phosphorylated at S396, we observed protein bands of the apparent molecular weight of 50-65 kDa in samples from both treated (OA+) and untreated (OA-) cells (**Fig. 1A**). Additionally, a protein co-migrating with the molecular weight of ∼100 kDa appeared in samples from cells treated with OA.

The same samples were analyzed with antibody CP13 that recognizes tau phosphorylated at S202 (**Fig. 1B**). In cells treated with OA a protein with an apparent molecular weight of ∼100 kDa was clearly visible, while in samples from untreated cells the signal was very weak or not detectable, indicating that under normal growth conditions endogenous tau is phosphorylated at S202 at low levels. A similar pattern of OA-induced 100 kDa phospho-tau immunoreactive protein was observed in SH-SY5Y cells differentiated into neuron-like cells (**Fig. 2**). Together, the data indicated that protein phosphatase inhibition by OA induced the formation of 100 kDa protein species immunoreactive to tau-pS202 and tau-pS396. Based on the molecular size of the reported protein and its immunoreactivity with anti-tau antibodies, this result is consistent with the possibility of tau dimerization.

**Figure 2.**
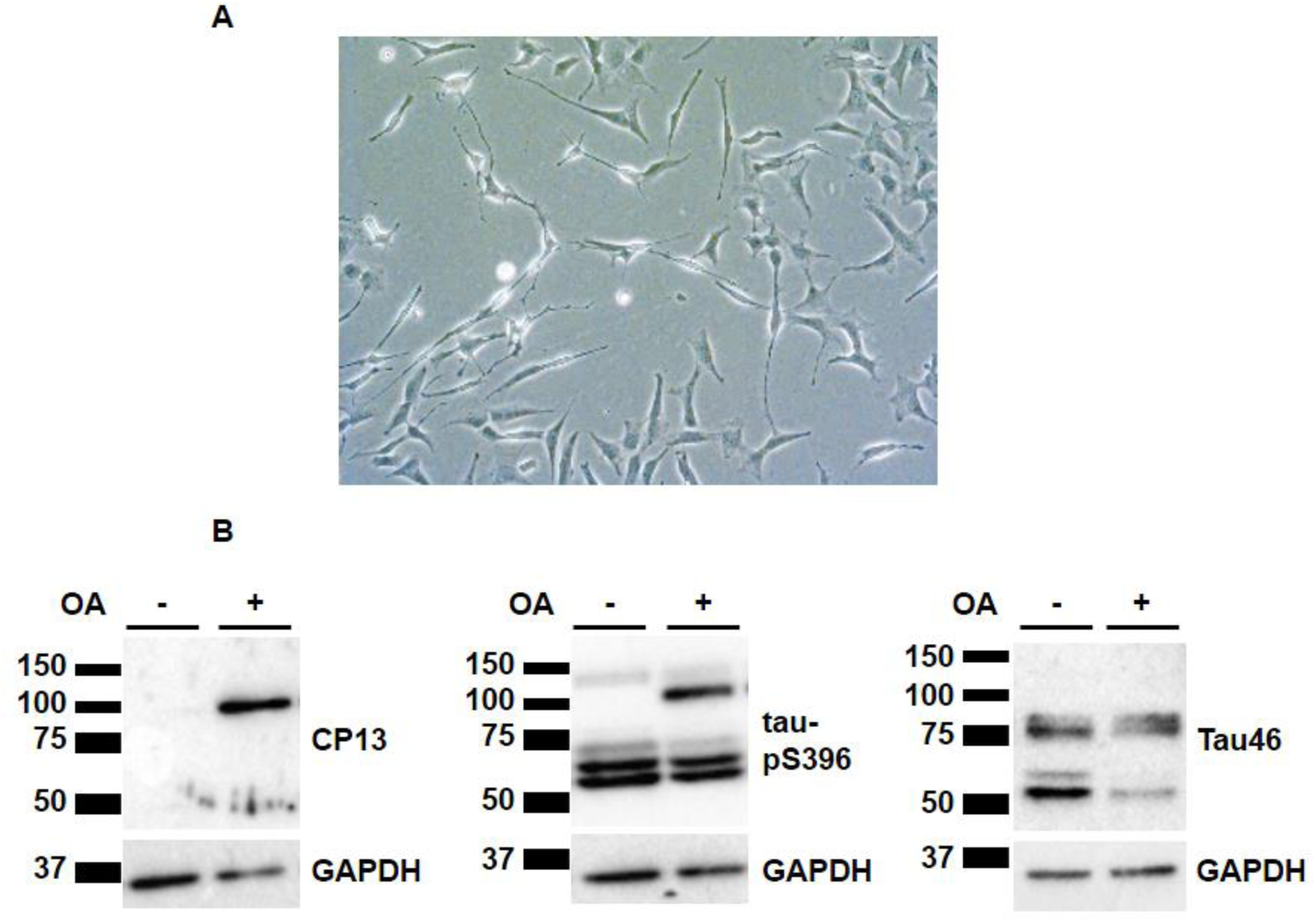
HMW-TIP is detected in OA-treated SH-SY5Y cells differentiated into neuron-like cells. Differentiated SH-SY5Y cells visualized using phase contrast microscopy (A) were treated with 100 nM OA for two hours. Note the differentiated cells with visible neuritic processes. Total cell lysates were analyzed by immunoblot using anti-tau-pS202 (CP13), anti-tau-pS396 and anti-total tau (Tau46) antibodies (B). Note a strong band migrating around 100-kDa in lysates of OA-treated cells. GAPDH is used as a loading control.

This high molecular weight tau-immunoreactive protein species (HMW-TIP) detected with CP13 and anti-tau-S396 antibodies migrated in the SDS-PAGE with nearly identical electrophoretic mobility. To test whether both CP13 and anti-tau-pS396 recognized the same protein, we divided the lane containing protein lysate from OA-treated cells on the nitrocellulose membrane in half by cutting it vertically. We incubated one half with antibody CP13 and the other half with antibody against tau-pS396 and aligned the halves for imaging (**Fig. 1C**). We noticed that the protein band recognized by the CP13 antibody migrated slightly slower than the anti-tau-pS396 antibody-reactive band (**Fig. 1C**), indicating two distinct protein species. These distinct protein bands could represent an identical tau splicing isoform containing different phosphorylation patterns or an additional posttranslational modification, such as protein truncation.

Next, we examined HMW-TIP using anti-total tau antibodies that recognize tau regardless of its phosphorylation state. We analyzed OA-treated and -untreated samples using antibody Tau46 that recognizes an epitope in the C-terminal part of tau (aa 404-441). We observed that antibody Tau46 did not detect the 100 kDa protein, although it did recognize 50 kDa tau and a protein around 70 kDa (**Fig. 3A and Fig. 2B**), which presumably represents MAP2. We further used anti-total tau antibodies raised against other tau regions, Tau13 (epitope most likely within the first 35 aa) and CP27 (whose epitope includes aa 130-150). Despite the recognition of monomeric tau proteins, Tau13 and CP27 antibodies did not detect HMW-TIP (**Figs. 3B and 3C**).

**Figure 3.**
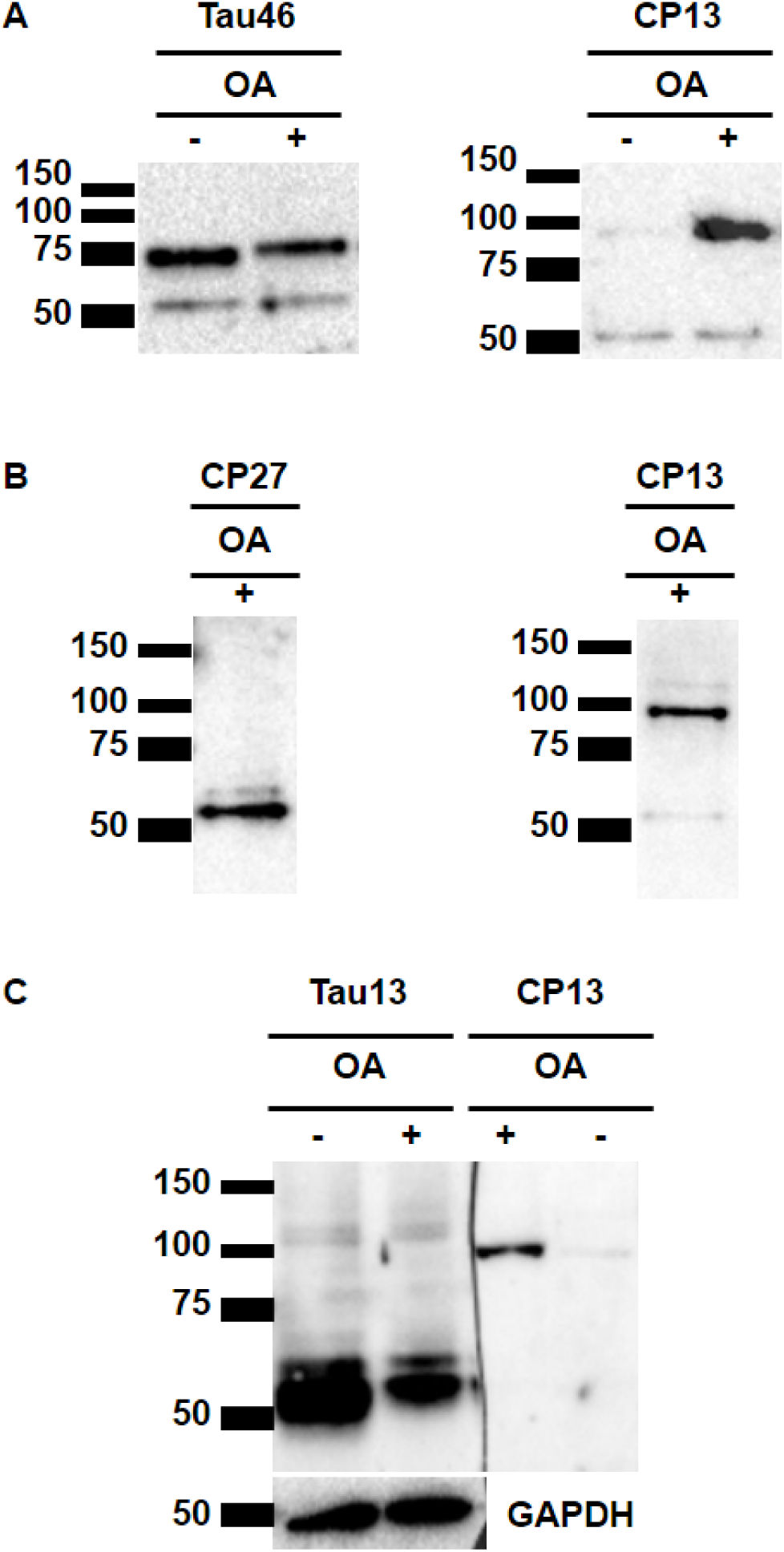
HMW-TIPs is not detected by anti-total tau antibodies. Where indicated, cells were treated with 100 nM (A) or 150 nM (B, C) OA+ for two hours. Total cells lysates were analyzed by immunoblotting using anti-total tau antibodies Tau46 (A), CP27 (B) and Tau13 (C). To visualize HMW-TIP, same samples were analyzed by antibody CP13. Note a CP13-reactive band migrating around 100-kDa in lysates of OA-treated cells (A, B, C). Note that all three anti-total antibodies detect a band migrating above 50 kDa that represents monomeric tau, but do not detect 100 kDa protein. Tau46 antibody additionally detects a 70 kDa band that represents MAP2c (A). GAPDH served as a loading control (C).

Although protein phosphatase inhibition by OA resulted in the detection of 100 kDa proteins immunoreactive to CP13 and anti-tau-pS396, we could not detect HMW-TIP using anti-total tau antibodies Tau13, CP27, and Tau46, which together cover tau epitopes in the N-terminal, central and C-terminal regions of tau. The fact that HMW-TIP was not detected using these antibodies could be due to antigen masking within the oligomer or a combination of epitope masking and protein truncation.

### 2.2. HMW-TIP is found in a heat-stable fraction

To investigate the properties of HMW-TIP, we examined whether it could be found in a heat-stable fraction. Due to their internally disordered structure, tau and other MAP2 family proteins are resistant to heat-induced precipitation. After isolating a heat-stable fraction using the protocol described by Petry and collaborators [19], in which cells are lysed in a buffer containing 1% non-ionic detergent Triton X-100, 0.25 % ionic detergent sodium-deoxycholate and 150 mM NaCl, but no SDS, the cell lysate was boiled and insoluble proteins were precipitated by centrifugation at 20,000 g. As expected, antibody Tau46 detected 50 kDa tau and a 70 kDa protein, presumably MAP2C in the heat-stable fraction, and no signal was observed in the pellet fraction (**Fig. 4A**, upper panel). In contrast, GAPDH levels were much lower in the heat-stable fraction than in the pellet (**Fig. 4A**, lower panel), indicating that the heat-stable fraction was enriched in heat-stable proteins. Using the antibody anti-tau-pS396, we found that HMW-TIP was present in the heat-stable fraction (**Fig. 4B**, left panel, HS), while it was undetectable in the pellet (**Fig. 4B**, right panel), similar to the distribution of monomeric tau detected by Tau46 (**Fig. 4A**). In conclusion, HMW-TIP co-fractionated with heat-stable proteins, which is a characteristic of proteins with internally disordered structure, including tau.

**Figure 4.**
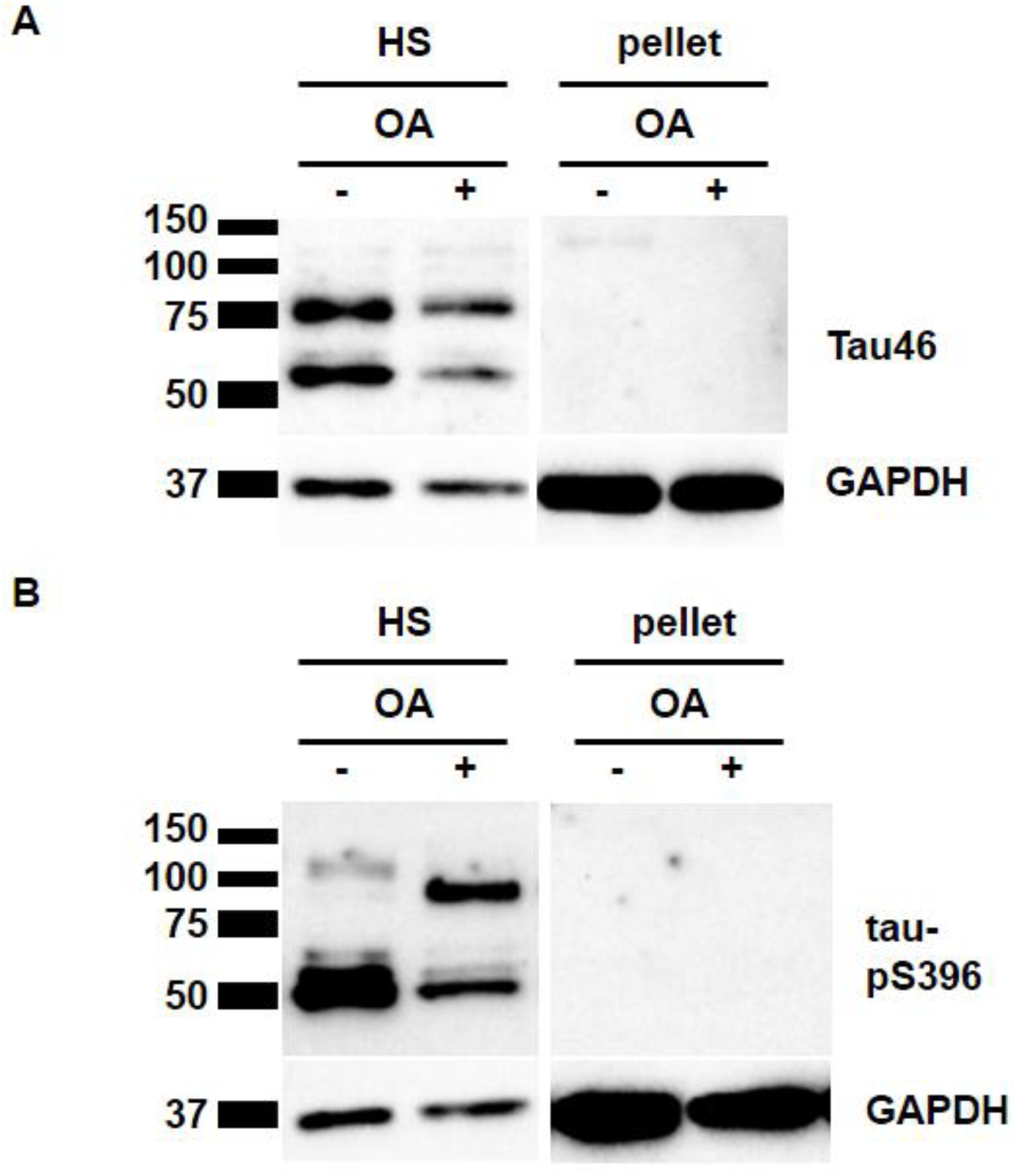
HMW-TIP is present in heat-stable fraction. SH-SY5Y cells differentiated into neuron-like cells were treated with 100 nM OA+ or DMSO (OA-) for two hours. Heat-stable (HS) and pellet fractions were analyzed by immunoblot using anti-total tau antibody Tau46 (A), anti-tau-pS396 (B) and anti-GAPDH (A, B) antibodies. Heat-stable fraction and pellet samples were analyzed on the same nitrocellulose membrane and imaged simultaneously. Note that 50 kDa tau and 70 kDa MAP2c detected by Tau46 antibody are enriched in the heat-stable fraction compared to pellet (A), similarly to 50 kDa tau and 100 kDa protein detected by anti-tau-pS396 (B). In contrast, GAPDH is enriched in pellet compared to heat-stable fraction (A, B).

### 2.3. HMW-TIP is stable under reducing and strong denaturing conditions

Considering the possibility that HMW-TIP represents a tau oligomer, we tested conditions under which oligomers could be dissociated. First, we compared HMW-TIP protein levels from cell lysates prepared under reducing and non-reducing conditions. Previous reports have shown that tau oligomers can be disrupted under reducing conditions (Sahara *et al.*, 2007, 2013), although tau crosslinking by disulfide bonds is not an absolute prerequisite for tau aggregation. As shown in the **Figure 5A**, CP13-immunoreactive HMW-TIP was similarly present in extracts prepared both in absence and presence of 5% β-mercaptoethanol, indicating that the stability of putative oligomer does not depend on disulfide bonds. Next, we examined whether putative oligomers could be dissociated in the presence of strong denaturing agent urea and guanidine hydrochloride. We lysed OA-treated cells in the lysis buffer containing 8 M urea or 6 M guanidine hydrochloride and analyzed total cell extracts by immunoblot. As shown in **Figures 5B and 5C**, HMW-TIP did not dissociate in the presence of either 8 M urea (**Fig. 5B**) or 6 M guanidine hydrochloride (**Fig. 5C**).

**Figure 5.**
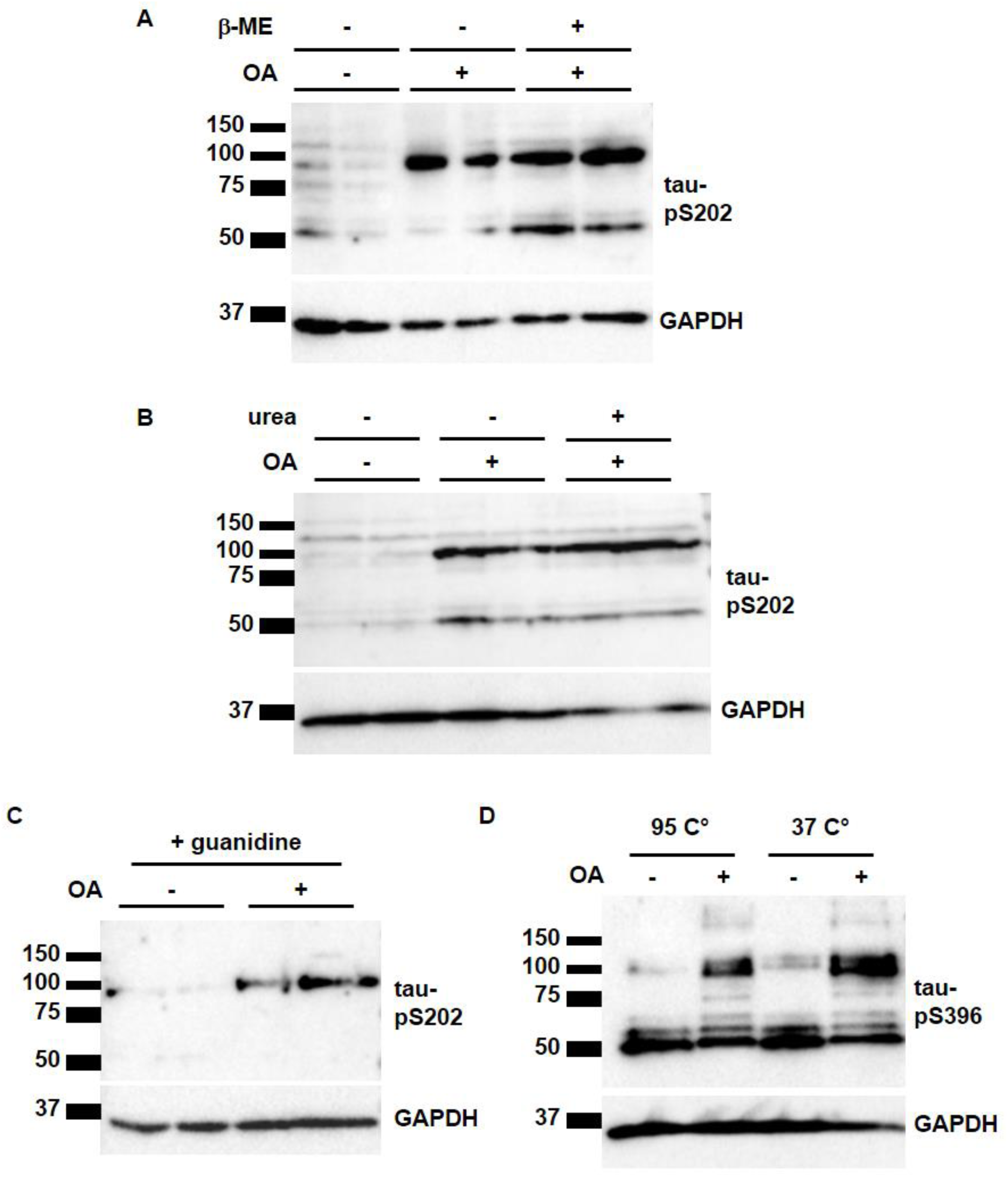
HMW-TIP stability under reducing and denaturing conditions. Cells were treated with 150 nM (A - C) or 100 nM (D) OA. Cell lysates were denatured at 37°C or 95°C (A). Cell lysates were prepared in absence or presence of 5 % β-mercaptoethanol (B), 8 M urea (C) or 6 M guanidine-hydrochloride (D). Immunoblotting was performed using anti-tau-pS202 (CP13) and anti-tau-pS396 antibodies. GAPDH was used as loading control. Note that the 100 kDa band is present in samples from OA-treated cells under all tested conditions.

As some proteins are prone to aggregation when subjected to denaturation at high temperature, we additionally tested whether the appearance of the 100 kDa protein could be a consequence of protein aggregation following denaturation of cell lysates at 95°C. We examined HMW-TIP levels in samples denatured at 37°C and 95°C and observed that they were similar (**Fig. 5D**), indicating that theHMW-TIP is not simply a consequence of protein aggregation due to a high temperature.

### 2.4. Characterization of the HMW-TIP species upon protein dephosphorylation

Finally, to examine the impact of protein phosphorylation on this putative 100 kDa tau oligomer, lysates of cells treated with OA were incubated with alkaline phosphatase. Because none of the tested anti-total tau antibodies recognized HMW-TIP, this presented an obstacle in examining proteins upon dephosphorylation. Surprisingly, we observed that alkaline phosphatase treatment did not abolish detection of 55-60 kDa tau and HMW-TIP by anti-tau-pS396 antibody (**Fig. 6**), although their electrophoretic mobility was slightly modified (**Fig. 6**, compare molecular weight of proteins in the samples treated and not treated with alkaline phosphatase). The change in electrophoretic mobility indicates that some amino acid residues were dephosphorylated, but not the S396 epitope. The fact that alkaline phosphatase treatment affects the migration rate of HMW-TIP only to a small degree suggested that the putative oligomer does not dissociate into monomers upon partial protein dephosphorylation.

**Figure 6.**
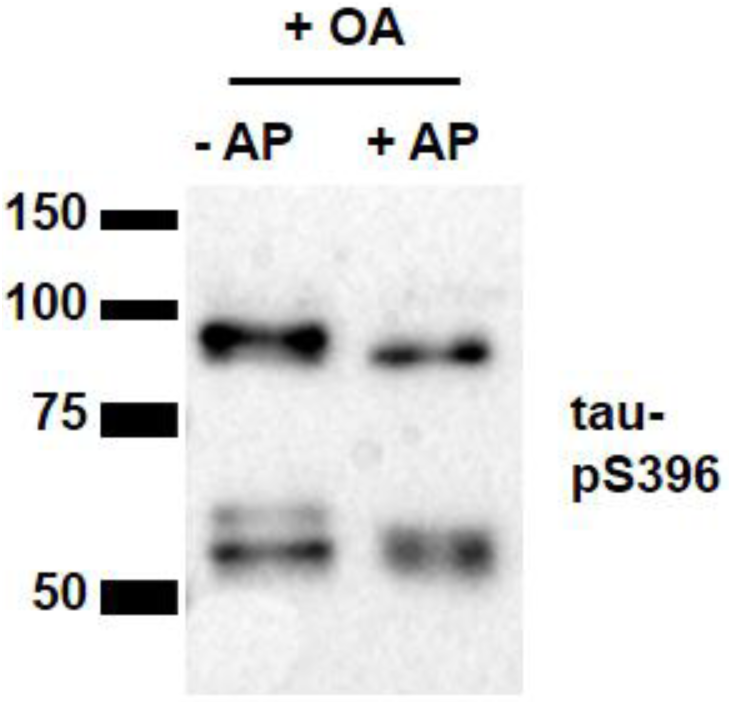
HMW-TIP stability upon protein dephosphorylation. Cells were incubated with 100 nM okadaic acid (OA) for two hours. Heat-stable fraction was treated with alkaline phosphatase and analyzed by immunoblot using anti-tau-pS396 antibody. Note the persistent antibody detection and changed electrophoretic mobility of proteins treated with alkaline phosphatase.

## 3. Discussion

In the present study we showed that inhibition of protein phosphatases by treatment of SH-SY5Y neuroblastoma cells with OA leads to the appearance of 100 kDa protein species immunoreactive to phospho-tau specific antibodies CP13 (anti-tau-pS202) and anti-tau-pS396.

Based on its molecular size, we initially considered the possibility that HMW-TIP represents a large tau splicing isoform containing exon 4A (so-called “big tau”), which contains an additional 254 aa insert in the N-terminal part [20] and is normally expressed predominantly in the peripheral nervous tissue [21]. This would be in accordance with a previous report showing that SH-SY5Y cells and their parental line SK-N-SH express a tau isoform around 100 kDa [15]. However, we were unable to detect HMW-TIP using any of the antibodies used against total tau, namely Tau46, CP27, and Tau13. Moreover, antibodies Tau5 and a rabbit polyclonal antibody also did not recognize HMW-TIP in our experiments. As “big tau” normally contains all of the epitopes recognized by these antibodies, with the possible exception of CP27, and these epitopes are not recognized by the anti-total tau antibodies we used, it is highly unlikely that HMW-TIP represents “big tau”.

Based on the observation that the molecular weight of the HMW-TIP corresponds to the predicted size of two molecules of fetal tau, which is one of the most prominent tau isoforms in undifferentiated SH-SY5Y cells [16], we considered the possibility that it represents tau oligomers. In support of this possibility, a study using bimolecular fluorescence complementation showed that HEK293 cells expressing tau constructs fused to split Venus increase in fluorescence upon treatment of cells with 30 nM OA for 24 h, suggesting tau oligomerization [22]. The absence of oligomer detection by anti-total tau antibodies in our experiments could be a consequence of by epitope masking or a combination of epitope masking and protein truncation, which would render HMW-TIP unrecognizable.

In attempt to dissociate putative oligomers by reducing and denaturing agents, we found that HMW-TIP was stable in the presence of β-mercaptoethanol, SDS, urea and guanidine. The role of disulfide bridges in tau oligomerization is not entirely clear, as both reduction-sensitive and reduction-resistant tau dimers have been previously observed [23, 24]. In our experiment β-mercaptoethanol did not eliminate the HMW-TIP signal, indicating that the stability of a putative dimer does not depend on disulfide bonds. Despite the fact that most protein complexes dissociate in the presence of SDS, some protein complexes remain assembled [25]. For example, SDS-resistant tau oligomers have been reported in AD brain [23]. Moreover, it has been previously reported that a pool of tau isolated from AD brain remained aggregated in high molecular mass structures even after the treatment with 8 M urea [26], whereas certain types of tau aggregates were stable in 6 M guanidine hydrochloride [27]. Taken together, we were unable to dissociate putative oligomers with SDS, urea, and guanidine. However, the observed stability of HMW-TIP in the presence of these agents is compatible with the characteristics of some previously reported tau oligomers.

The present study showed that treatment of cell lysates with alkaline phosphatase resulted in partially dephosphorylated HMW-TIP, leaving the S396 epitope phosphorylated and the putative oligomer assembled. In fact, an inefficient removal of phosphates by alkaline phosphatase from the tau PHF-1 epitope, which includes pS396 and pS404, has been already reported [28]. Our data suggest that exhaustive tau phosphorylation is either not essential for the stability of the putative oligomer, or is necessary for initial oligomer formation, but once formed, oligomers could be stable even upon dephosphorylation. Based on our present data it is also possible that, in addition to other sites, dephosphorylation of S396 is required for the disassembly of the putative oligomer.

In summary, protein phosphatase inhibition by OA lead to appearance of HMW proteins immunoreactive to anti-tau-pS202 and -pS396 antibodies, which is in line with the possibility that HMW-TIP represent tau oligomers. However, due to the inability to detect HMW-TIP with anti-total tau antibodies and to dissociate putative oligomers under conditions tested, we cannot exclude the possibility that HMW-TIP represent tau-unrelated proteins that are detected with anti-tau-pS202 and -pS396 antibodies due to cross-reactivity. The finding that anti-tau-pS396- and -pS202-reactive HMW-TIP exhibited almost identical, yet distinct electrophoretic mobilities in SDS-PAGE supports this possibility. Furthermore, although HMW-TIP was present in the heat-stable fraction, which is a characteristic of tau and other intrinsically disordered proteins [29], it should be noted that due to the presence of 1% Triton X-100 and 0.25 % sodium-deoxycholate in the lysis buffer, our protocol may have extracted a broader range of proteins than buffers without detergents.

In conclusion, we provide convincing evidence that OA treatment induced HMW phospho-tau immunoreactive proteins in a neuroblastoma cell culture. Although further research is required to clarify the identity of the reported proteins, these findings will facilitate future studies of tau protein related processes under conditions of protein phosphatase inhibition.

## Acknowledgments

We would like to thank Dr. Peter Davies (Albert Einstein College of Medicine, Bronx, NY, USA) for the gift of CP13 and CP27 antibodies.

## Author Contributions

### Declaration of authorship

GŠ conceived and directed the project, coordinated experiments and co-wrote the paper. MB designed, performed and interpreted experiments and co-wrote the paper. TM performed and analyzed experiments under supervision of MB. MBL established the experimental model and performed and interpreted initial experiments. PRH substantially contributed to the interpretation of data for the work. All authors contributed to revising and editing the manuscript critically for important intellectual content. All authors approved the final version of the manuscript.

## Abbreviations

AD: Alzheimer’s disease
BDNF: brain-derived neurotrophic factor
CP13: antibody against phosphorylated tau Ser-202
CP27: antibody against total tau (aa 130-150)
DMEM: Dulbecco’s modified Eagle medium
DMSO: dimethyl sulfoxide
EDTA: ethylenediaminetetraacetic acid
FBS: fetal bovine serum
GAPDH: glyceraldehyde 3-phosphate dehydrogenase
GSK-3β: glycogen synthase kinase 3β
HMW-TIP: high molecular weight tau-immunoreactive protein
HS: heat-stable (fraction)
MAP: microtubule-associated protein
MTB: microtubule binding (repeat regions)
NFT: neurofibrillary tangles
OA: okadaic acid
PBS: phosphate-buffered saline
PP2A: protein phosphatase 2A
RIPA: radioimmunoprecipitation assay buffer
SDS-PAGE: sodium dodecyl sulfate – polyacrylamide gel electrophoresis
Tau5: antibody against the central part of tau (aa 210-241)
Tau13: antibody against the N-terminal part of tau (aa 2-18)
Tau46: antibody against the C-terminal part of tau (aa 404-441)
TBS: Tris-buffered saline

## References

1. Binder, L.I.; Frankfurter, A.; Rebhun, L.I. The distribution of tau in the mammalian central nervous system. J. Cell Biol. 1985, 101, 1371–1378.

2. Buée, L.; Bussière, T.; Buée-Scherrer, V.; Delacourte, A.; Hof, P.R. Tau protein isoforms, phosphorylation and role in neurodegenerative disorders. Brain Res. Rev. 2000, 33, 95–130.

3. Šimić, G.; Babić Leko, M.; Wray, S.; Harrington, C.; Delalle, I.; Jovanov-Milošević, N.; Bažadona, D.; de Silva, R.; Di Giovanni, G.; Wischik, C; Hof, P.R. Tau protein hyperphosphorylation and aggregation in Alzheimer’s disease and other tauopathies, and possible neuroprotective strategies. Biomolecules 2016, E6.

4. Uversky, V.N. Natively unfolded proteins: a point where biology waits for physics. Protein Sci. 2002, 11, 739–756.

5. Zhou, Y.; Shi, J.; Chu, D.; Hu, W.; Guan, Z.; Gong, C.X.; Iqbal, K.; Liu, F. Relevance of phosphorylation and truncation of tau to the etiopathogenesis of Alzheimer’s disease. Front. Aging Neurosci. 2018, 10, 27.

6. Jeganathan, S.; von Bergen, M.; Brutlach, H.; Steinhoff, H.J.; Mandelkow, E.; Global hairpin folding of tau in solution. Biochemistry 2006, 45, 2283–2293.

7. Wang, Y.; Mandelkow, E. Tau in physiology and pathology. Nat. Rev. Neurosci. 2016, 17, 22–35.

8. Duka, V.; Lee J.H.; Credle, J.; Wills, J.; Oaks, A.; Smolinsky, C.; Shah, K.; Mash, D.C.; Masliah, E.; Sidhu, A. Identification of the sites of tau hyperphosphorylation and activation of tau kinases in synucleinopathies and Alzheimer’s diseases. PLoS One 2013, 8, e75025.

9. Martin, L.; Latypova, X.; Wilson, C.M.; Magnaudeix, A.; Perrin, M.L.; Terro, F. Tau protein phosphatases in Alzheimer’s disease: the leading role of PP2A. Ageing Res. Rev. 2013, 12, 39–49.

10. Liu, F.; Grundke-Iqbal, I.; Iqbal, K.; Gong, C.X. Contributions of protein phosphatases PP1, PP2A, PP2B and PP5 to the regulation of tau phosphorylation. Eur. J. Neurosci. 2005, 22, 1942–1950.

11. Bennecib, M.; Gong, C.X.; Grundke-Iqbal, I.; Iqbal, K. Role of protein phosphatase-2A and −1 in the regulation of GSK-3, cdk5 and cdc2 and the phosphorylation of tau in rat forebrain. FEBS Lett. 2000, 485, 87–93.

12. Arif, M.; Wei, J.; Zhang, Q.; Liu, F.; Basurto-Islas, G.; Grundke-Iqbal, I.; Iqbal, K. Cytoplasmic retention of protein phosphatase 2A inhibitor 2 (I2PP2A) induces Alzheimer-like abnormal hyperphosphorylation of Tau. J. Biol. Chem. 2014, 289, 27677–27691.

13. Swingle, M.; Ni, L.; Honkanen, R.E. Small-molecule inhibitors of Ser/Thr protein phosphatases: specificity, use and common forms of abuse. Methods Mol. Biol. 2007, 365, 23–38.

14. Smith, C.J.; Anderton, B.H.; Davis, D.R.; Gallo, J.M. Tau isoform expression and phosphorylation state during differentiation of cultured neuronal cells. FEBS Lett. 1995, 375, 243–248.

15. Uberti, D.; Rizzini, C.; Spano, P.F.; Memo, M. Characterization of tau proteins in human neuroblastoma SH-SY5Y cell line. Neurosci. Lett. 1997, 235, 149–153.

16. Agholme, L.; Lindström, T.; Kågedal, K.; Marcusson, J.; Hallbeck, M. An in vitro model for neuroscience: differentiation of SH-SY5Y cells into cells with morphological and biochemical characteristics of mature neurons. J. Alzheimers Dis. 2010, 20, 1069–1082.

17. Encinas, M.; Iglesias, M.; Liu, Y.; Wang, H.; Muhaisen, A.; Ceña, V.; Gallego, C.; Comella, J.X. Sequential treatment of SH-SY5Y cells with retinoic acid and brain-derived neurotrophic factor gives rise to fully differentiated, neurotrophic factor-dependent, human neuron-like cells. J. Neurochem. 2000, 75, 991–1003.

18. Pepinsky, R.B. Selective precipitation of proteins from guanidine hydrochloride-containing solutions with ethanol. Anal. Biochem. 1991, 195, 177–181.

19. Petry, F.R.; Pelletier, J.; Bretteville, A.; Morin, F.; Calon, F.; Hébert, S.S.; Whittington, R.A.; Planel, E. Specificity of anti-tau antibodies when analyzing mice models of Alzheimer’s disease: problems and solutions. PLoS One 2014, 9, e94251.

20. Goedert, M.; Spillantini, M.G.; Crowther, R.A. Cloning of a big tau microtubule-associated protein characteristic of the peripheral nervous system. Proc. Natl. Acad. Sci. USA 1992, 89, 1983–1987.

21. Boyne, L.J.; Tessler, A.; Murray, M.; Fischer, I. Distribution of Big tau in the central nervous system of the adult and developing rat. J. Comp. Neurol. 1995, 358, 279–293.

22. Tak, H.; Haque, M.M.; Kim, M.J.; Lee, J.H.; Baik, J.H.; Kim, Y.; Kim, D.J.; Grailhe, R.; Kim, Y.K. Bimolecular fluorescence complementation; lighting-up tau-tau interaction in living cells. PLoS One 2013, 8, e81682.

23. Sahara, N.; Maeda, S.; Murayama, M.; Suzuki, T.; Dohmae, N.; Yen, S.H.; Takashima, A. Assembly of two distinct dimers and higher-order oligomers from full-length tau. Eur. J. Neurosci. 2007, 25, 3020–3029.

24. Sahara, N.; DeTure, M.; Ren, Y.; Ebrahim, A.S.; Kang, D.; Knight, J.; Volbracht, C.; Pedersen, J.T.; Dickson, D.W.; Yen, S.H.; Lewis, J. Characteristics of TBS-extractable hyperphosphorylated tau species: aggregation intermediates in rTg4510 mouse brain. J. Alzheimers Dis. 2013, 33, 249–263.

25. Bitan, G.; Fradinger, E.A.; Spring, S.M.; Teplow, D.B. Neurotoxic protein oligomers – what you see is not always what you get. Amyloid 2005, 12, 88–95.

26. Köpke E.; Tung, Y.C.; Shaikh, S.; Alonso, A.C.; Iqbal, K.; Grundke-Iqbal, I. Microtubule-associated protein tau. Abnormal phosphorylation of a non-paired helical filament pool in Alzheimer disease. J. Biol. Chem. 1993, 268, 24374–24384.

27. Falcon, B.; Cavallini, A.; Angers, R.; Glover, S.; Murray, T.K.; Barnham, L.; Jackson, S.; O’Neill, M.J.; Isaacs, A.M.; Hutton, M.L.; Szekeres, P.G.; Goedert, M.; Bose, S. Conformation determines the seeding potencies and recombinant Tau aggregates. J. Biol. Chem. 2015, 290, 1049–1065.

28. Tepper, K.; Biernat, J.; Kumar, S.; Wegmann, S.; Timm, T.; Hübschmann, S.; Redecke, L.; Mandelkow, E.M.; Müller, D.J.; Mandelkow, E. Oligomer formation of tau protein hyperphosphorylated in cells. J. Biol. Chem. 2014, 289, 34389–34407.

29. Battisti, A.; Ciasca, G.; Grottesi, A.; Tenenbaum, A. Thermal compaction of the intrinsically disordered protein tau: entropic, structural, and hydrophobic factors. Phys. Chem. Chem. Phys. 2017, 19, 8435–8446.

